# Distinct Functions for Beta and Alpha Bursts in Gating of Human Working Memory

**DOI:** 10.1101/2023.11.17.566386

**Authors:** Johan Liljefors, Rita Almeida, Gustaf Rane, Johan N. Lundström, Pawel Herman, Mikael Lundqvist

## Abstract

Multiple neural mechanisms underlying gating to and from working memory (WM) have been proposed, with divergent results obtained in human and animal studies. Previous results from non-human primate studies suggest information encoding and retrieval is regulated by high-power bursts in the beta frequency range, whereas human studies suggest that alpha power in sensory regions filters out unwanted stimuli from entering WM. Discrepancies between studies, whether due to differences in analysis, species, or cortical regions, remain unexplained. We addressed this by performing similar single-trial burst analysis we earlier deployed on non-human primates on human whole-brain electrophysiological activity. Participants performed a sequential working memory task that allowed us to track the distinct electrophysiological activity patterns associated with neural processing of targets and distractors. Intriguingly, our results reconcile earlier findings by demonstrating that both alpha and beta bursts are involved in the filtering and control of WM items, but with region and task-specific differences between the two rhythms. Occipital beta burst patterns regulate the transition from sensory processing to WM retention whereas prefrontal and parietal beta bursts track sequence order and proactively suppress retained information prior to upcoming target encoding. Occipital alpha bursts instead suppress unwanted sensory stimuli during their presentation. These results suggest that human working memory is regulated by multiple neural mechanisms that operate in different cortical regions and serve distinct computational roles.

## INTRODUCTION

Working memory (WM) is a key cognitive component that allows us to hold and manipulate information online in our mind (D’Esposito & Postle, 2015; Goldman−Rakic, 1995; Ma et al., 2014; Miller et al., 2018; Vogel et al., 2005). WM has a very limited capacity and thus, we need to control that only relevant information enters our WM (Chatham & Badre, 2015; Liesefeld et al., 2014; Vogel et al., 2005). Work in non-human primates and humans have, however, suggested distinct mechanisms of such WM control. Here, we resolve this discrepancy and demonstrate multiple novel task- and region-specific neural correlates of human WM control on a single trial level.

Using intracranial recordings in prefrontal cortex of non-human primates, we previously observed high power beta (20-35 Hz) bursts as a single-trial-correlate of executive control over WM (Bastos et al., 2018; Lundqvist et al., 2016, 2018, 2020, 2023). Specifically, beta bursting was reduced when information was encoded, particular in cortical sites in which neurons subsequently retained the information in WM (Lundqvist et al., 2016, 2018). This suggests that spatial patterns of prefrontal beta bursts filter out unwanted stimuli. Similarly, prefrontal beta bursts were suppressed at time points in which WM was accessed and subsequently elevated when WM was cleared out.

In contrast, investigations of human participants instead suggest alpha (8-12 Hz) power in occipital and parietal areas as a correlate of filtering out visual information (Gutteling et al., 2022; Popov et al., 2017, 2019; Roux & Uhlhaas, 2014; Sauseng et al., 2005; Turner et al., 2023). Alpha power is suppressed at cortical locations that process relevant information and is elevated at others. These spatio-temporal patterns of alpha are selective to the spatial location of distractors or their sensory modality (Haegens et al., 2012; Popov et al., 2019; Turner et al., 2023; Yuasa et al., 2023; Zhou et al., 2023). In addition, in analogy with prefrontal beta bursts in non-human primates, alpha oscillations in human occipital and parietal regions have an inhibitory role in other WM-related control processes such as removal and selective prioritization of information (Riddle et al., 2020; Wolff et al., 2017).

Thus, the control-related patterns we observed in beta bursting in frontal cortex of non-human primates appear to be analogous to the ones observed in parieto-occipital alpha power fluctuations in humans. This difference in frequency could be related to species differences, analysis methods (power fluctuations vs bursts) or the distinct areas studied. The observation that the frequency of prefrontal beta power and bursts are gradually shifted towards frequencies lower in the cortical hierarchy, and occurring in the alpha range in visual area V4, would suggest the latter (Hipp et al., 2012; Lundqvist et al., 2020; Rosanova et al., 2020).

To test this hypothesis in human participants, we deployed a sequential WM task that required both input and output gating. To determine the single trial neural correlates of WM gating, we recorded whole scalp magnetoencephalogram (MEG) and frequency tagged each WM item in the sequence (Gutteling et al., 2022; Parkkonen et al., 2008; Zhigalov et al., 2019). Frequency tagging in our task involves modulating the luminance of stimuli at a known frequency which entrains cerebral activity. This allowed us to estimate the degree individual items were processed depending on their status as a target or distractor, and how this related to alpha and beta bursting in different cortical regions. The goal was to reconcile the roles of beta bursting in macaques with alpha power in humans. Are both alpha and beta involved in the gating of distractor items? Do they have a role in gating information into working memory observable on single trials and in behaviour? If so, do they have distinct roles and cortical origins? We found evidence that high power bursts of both alpha and beta gate information in and out of working memory, but with partially distinct roles between the two frequency bands and between cortical locations. Overall, alpha seemed consistent with the suppression of sensory processing, whereas beta bursts more selectively appeared to gate information from sensory processing into WM and proactively removing information already retained in WM.

## RESULTS

Using MEG we recorded cerebral data from 17 healthy volunteers while they performed a serial WM task (Figure 1). The task required them to hold fixation on the centre of the screen throughout the whole trial. There were two sets of trials, with or without distractors. On No-Distractor trials, four randomly oriented bars were sequentially presented foveally for 500 ms with inter-stimulus periods of 500 ms showing a fixation dot. Each bar was associated with a colour. Following a delay of 750 ms the fixation dot turned into one of the four colours, acting as a retro-cue signalling which of the four bars would be tested. After an additional 750 ms, a test probe appeared on the screen and subjects were to orient the test probe using a control pad, to match the memorized and cued bar within 5 seconds. On Distractor trials, the second and third bars acted as distractors, not to be remembered and were never probed. Distractor and No-Distractor trials were randomly interleaved and identified by a pre-cue just prior to each trial (Figure 1).

**Figure 1.**
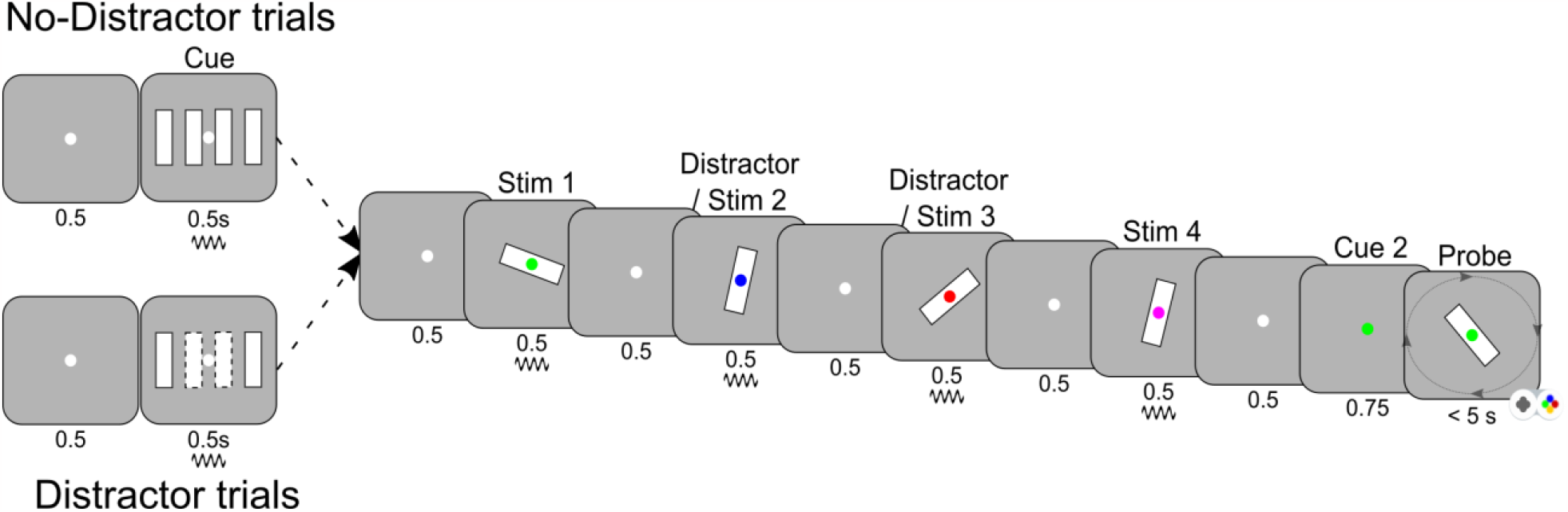
Experimental design. A sequential array of four bars with random orientation, was presented. Each bar was randomly associated with a colour (red, green, blue or magenta). Later, a retro-cue with one of the four colours was presented, identifying the target. In half the trials (randomized), bar 2 and 3 in the sequence were never tested and acted as distractors. These trials were indicated by a pre-cue. The pre-cue and all bars were frequency tagged (Methods). The subjects had 5 seconds to submit a response before the next trial started. Each subject performed 400 trials.

### Behavioural findings

To establish that subjects correctly performed the task and selectively encoded target bars, we first analysed behaviour. We measured performance as the absolute value of the angle between presented and reported orientation of the bar. The performance data was transformed in order to meet the general linear model assumptions (see Methods). The data was modelled using a linear mixed effects model, with subject as random effect. Condition (Distractor or No-Distractor), order of the bar probed and the interaction between these two variables were modelled as fixed effects. In the first model, trials probing bar 2 and 3 in the No-Distractor condition were removed to match the trials of the Distractor condition. Effect of condition was significant (t = 9.89, p < 2e-16), as well as order (t = -10.0, p < 2e-16). Further, the interaction between condition and bar probed was significant (p < 2e-7), meaning that the effect of condition was not the same for bar 1 and bar 4. The performance on the Distractor condition was better than on the No-Distractor condition, although the task difference for bar 1 (p<0.0001) was larger than for bar 4 (p=0.01). This suggested that participants indeed treated the second and third bar as distractors on these trials, freeing up resources to encode the two target bars on Distractor trials with higher precision.

Separating the data according to the condition, we found a clear serial order effect for both conditions, with much smaller errors on the last bar in the sequence relative to the first bar (p<0.0001). Finally, we analysed performance on the No-Distractor condition in relation to the bar probed, considering all four bars. There was a significant effect of bar (p<0.0001). Planned comparisons using Tukey’s method showed that the effect was driven by better performance on the last bar when compared to the three first ones, for which there were no statistically significant differences in performance (Figure 2).

**Figure 2.**
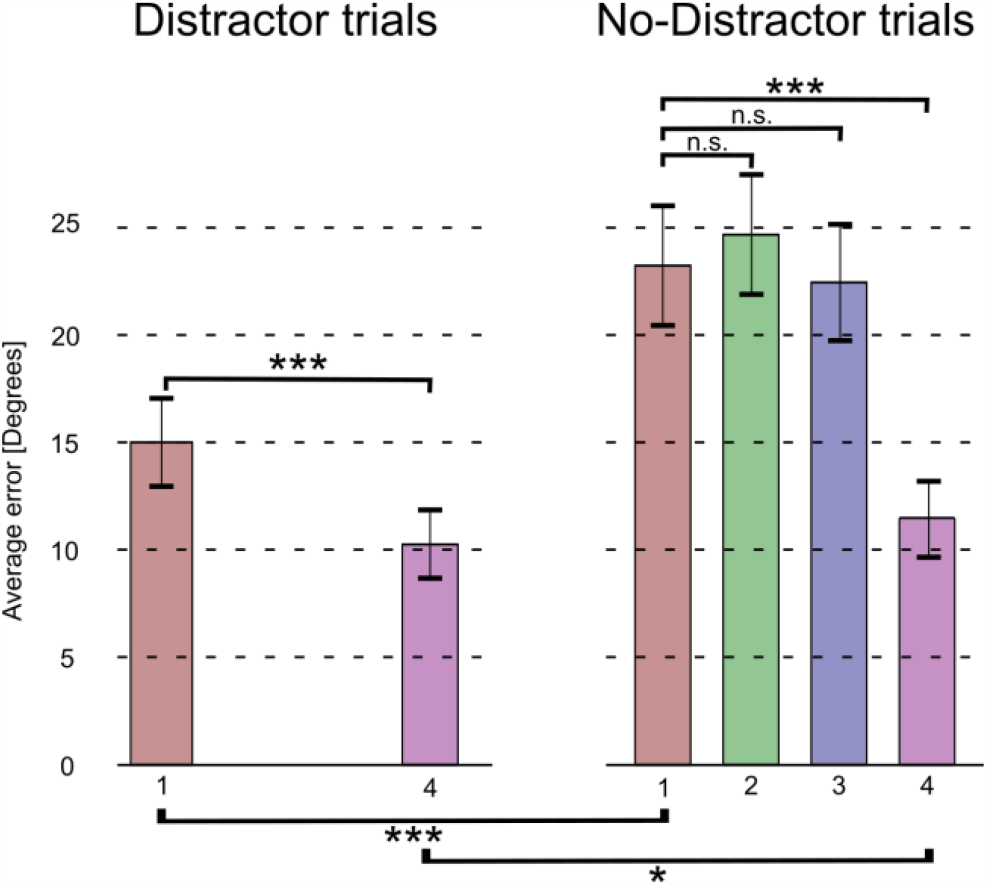
Performance by trial type. Average absolute error (angular distance between target and response) are shown with their ±1 SEM, for each bar in the sequence and by trial type (left Distractor trials, right No-Distractor trials). Significance was calculated by fitting linear mixed-effects models, and planned comparisons using Tukey’s method (see Methods).

Taken together, the results suggest that subjects appeared to encode all target bars and selectively skipped to encode distractors. The encoding of additional bars in the sequence degraded the memory representations of the bars already encoded, leading to a strong order effect that was more enhanced on trials in which more items were to be encoded. Thus, on the last bar in the sequence there was relatively good performance regardless of trial type.

### Frequency tagging revealed correlates of selective encoding

We next analysed the MEG signals at the sensor level. Each oriented bar as well as the pre-cue were frequency tagged (alternating between 31.1 or 37.1Hz, see Methods). Because the frequency tagging was phase-locked to the stimulus onset, we analysed total, phase-locked, and induced power separately. Neural activity corresponding to processing of the tagged stimuli would be more strongly observed in the phase-locked power as the tagging itself was phase-locked to the onset of stimulus (Figure 3A, right). We assessed neural substrates of gating by contrasting power in the No-Distractor and Distractor trials. This analysis demonstrated that distractors indeed entrained cortical activity to a lesser degree than targets (Figure 3A, right). It also revealed differences in induced power in the alpha and beta frequency ranges between these two conditions around the same time (Figure 3A, middle).

**Figure 3.**
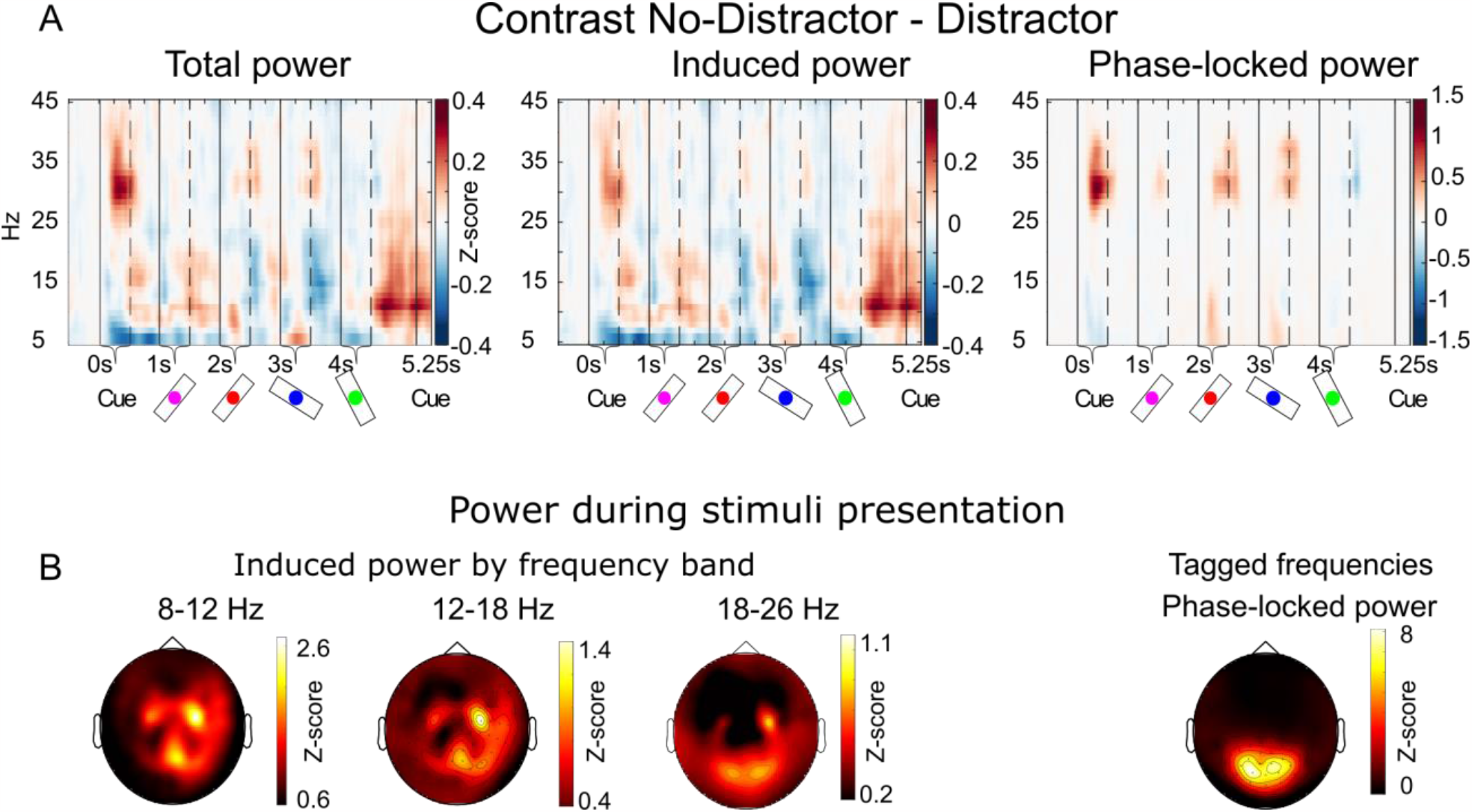
Time-frequency analysis of Distractor and No-Distractor trials. A) Power contrasts between the Distractor and No-Distractor trials. Spectral decompositions were done using adaptive superlets and z-scores were calculated on a trial level normalized to a pre-stimulus window of 250 ms before the onset of the cue (see Methods). Induced power was calculated by subtracting the event related field from the raw MEG data before spectral decomposition, and phase-locked power as the difference between total and induced power. Vertical lines show the onset/offset of the cues and bars. B) The topographical plots show the average induced power (z-score) for the three frequency bands, and the phase-locked power for the tagged frequencies (31.1Hz and 37.1 Hz).

### Both beta and alpha bursts selectively suppressed distractors

To directly connect our prior work on beta bursts in non-human primates to the present results, we extracted bursts in three frequency ranges (Alpha: 8-12 Hz; Low beta: 12-18 Hz; High beta: 18-26 Hz) for each sensor (Methods). The average burst rates, for all sensors and all trials, for both alpha and beta frequencies showed strong similarities with prefrontal beta burst rates in non-human primates (Figure 4A-B). Namely, the burst rates were elevated during fixation and delays, and suppressed during stimulus presentations. This is generally consistent with their proposed inhibitory role in filtering cortical bottom up processing (Lundqvist et al., 2016, 2018). There were, however, two striking differences between the alpha and beta burst rates over time. First, alpha peaked just before stimulus onset, was suppressed during each stimulus presentation but then started to smoothly rebound and peaking just before the next presentation. This rebound started even before the stimulus was removed. In beta, the pattern was similar but whereas alpha burst rates reacted similar to all stimuli in the sequence, beta burst rates were gradually lower as the sequence progressed (Figure 4B). We have previously observed the same for beta bursting in prefrontal cortex of non-human primates (Figure 6 in (Lundqvist et al., 2016)). Second, in contrast to alpha, beta burst rates were timed to both stimulus onset and offset, not just onset (Figure 4D). Both these differences were clearer in higher beta band compared to lower beta band, gradually changing from alpha to higher beta.

**Figure 4.**
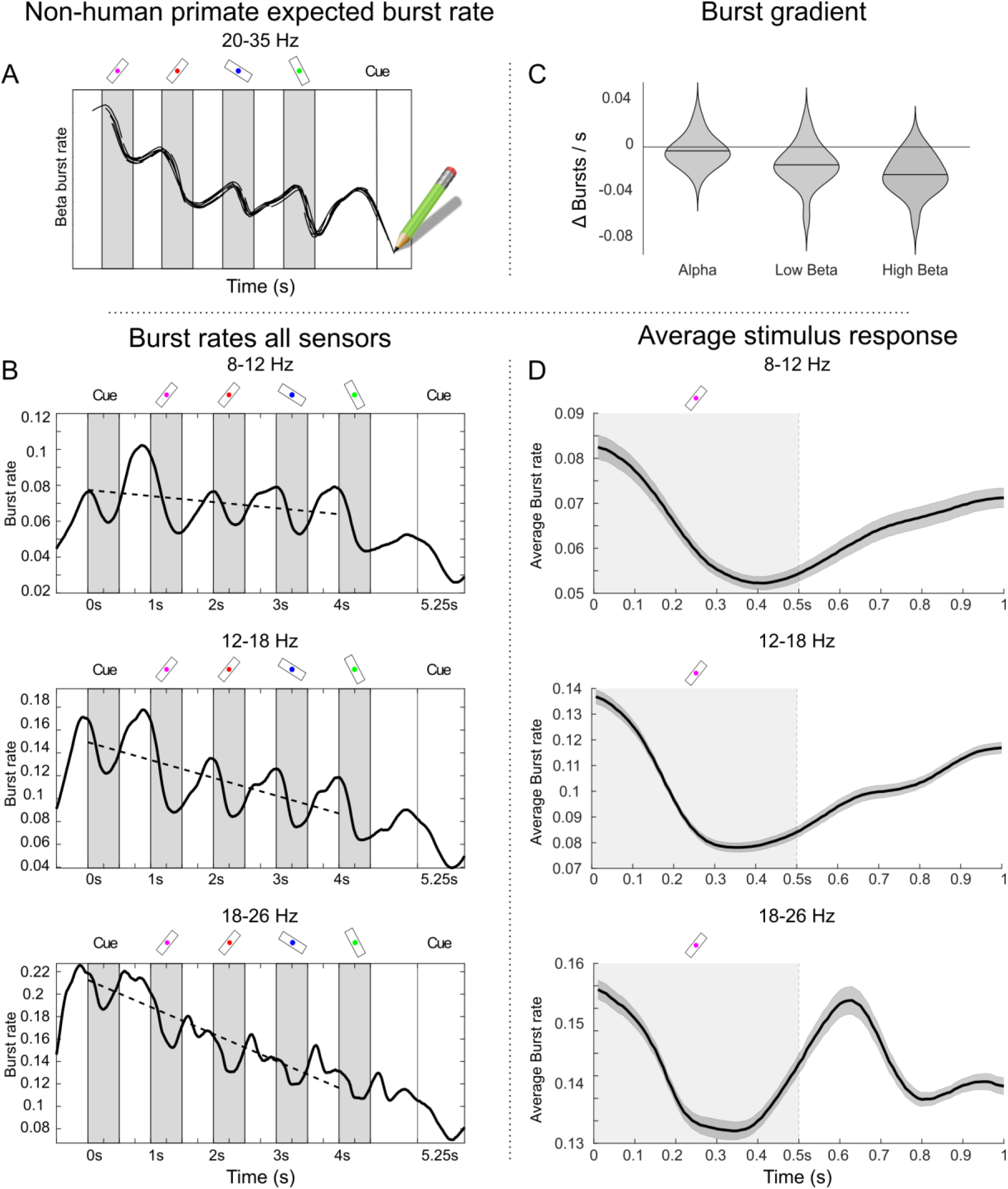
Burst rates across all sensors. A) Expected beta burst rate in Macaque monkeys. Based on prior research were monkeys were tasked to memorize 3 stimuli for periods of 0.3s (Lundqvist et al., 2016). The illustration has been adapted to be comparable to our experiment. B) Distinct temporal dynamics of alpha and beta bursts across all sensors. Plots show the grand average burst rates (across all sensors, trial types and subjects) for the three frequency bands. Burst rate is calculated as the fraction of trials where a burst was detected at that given part of the trial (Methods). The greyed areas represent the bar presentation, while the dotted line indicates the burst gradient, calculated as a linear fit using least squares. C) Burst gradient by frequency band for all trial types, represented as the slope of the dotted lines in figure 4B. Violin plots are based on subject level averages, horizontal line indicates mean. D) Burst rate from C), averaged over 1 second periods, across the 4 bars and their following delay period. Grey areas show stimulus presentation period.

### Alpha and beta bursts had distinct function depending on cortical origin

To better understand if the burst patterns observed in the alpha and beta bands were meaningful in terms of WM filtering, we next investigated how they differed in occipital, parietal and prefrontal regions and between Distractor and No-Distractor trials (Figure 5). There were several significant differences between the two conditions. First, there was increased beta and alpha bursting during and immediately following presentations of distractors as compared to targets (Figure 5). This difference was only seen in occipital sensors (see Figure 5 for all statistical differences referred to in this paragraph) and coincided with the decreased processing of distractors as measured by the phase-locked power in the tagged frequencies over occipital sensors (Figure 3). This elevated bursting during distractors was seen in all three frequency bands in occipital sensors, but had different characteristics depending on frequency (Figure 6). Alpha bursting was less suppressed during and following the presentation of targets relative distractors. Beta bursting, by contrast, had a distinct peak at around the time when distractor presentations ended (Figure 5, 6). This was consistent with their distinct temporal dynamics observed above, that beta (high beta in particular) appeared timed to both the onset and the offset of stimuli and not just onset as in the case of alpha. Thus, the difference between distractors and targets in beta bursting was more focused around the offset of the distractor, in the transition from sensory to working memory processing.

**Figure 5.**
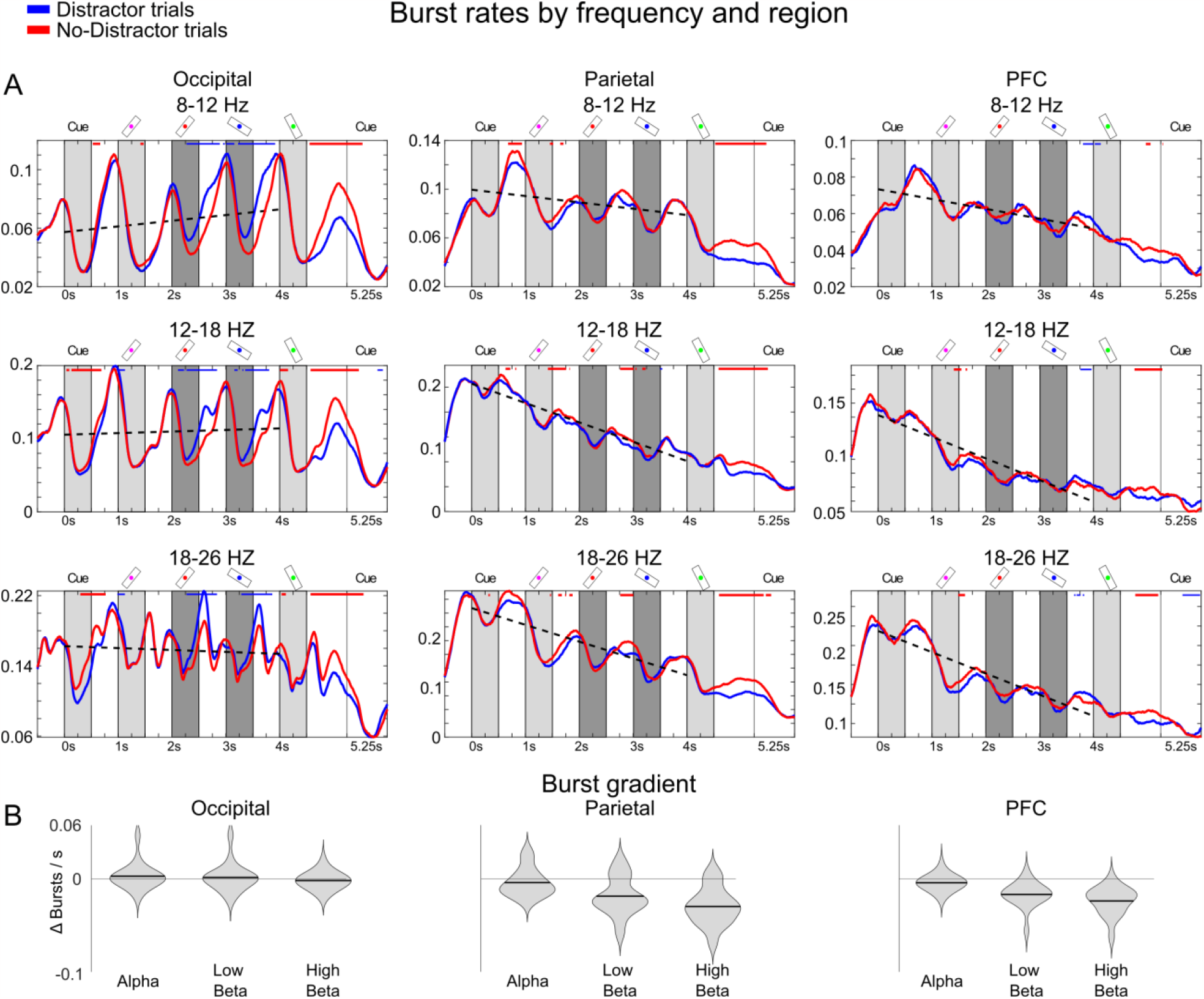
Bursting by cortical region. A) Burst rates differences between trial types and regions. Burst rates per frequency band (rows) and region (columns) are shown. Burst rates for Distractor (blue) and No-Distractor (red) trials are plotted independently. Blue bars denote when Distractor trial burst rates were significantly above No-Distractor trial burst rates, and red bars the opposite, using permutation tests at the p<0.001 level. The dotted line indicates the burst gradient, calculated through a linear fit using least squares to all trials. Potential distractor presentation periods (bar 2 and bar 3) are shown in dark (as opposed to light) grey areas. B) Burst gradient by frequency band and region for all trial types, represented as the slope of the dotted lines in A). Violin plots are based on subject level averages, horizontal line indicates mean.

**Figure 6.**
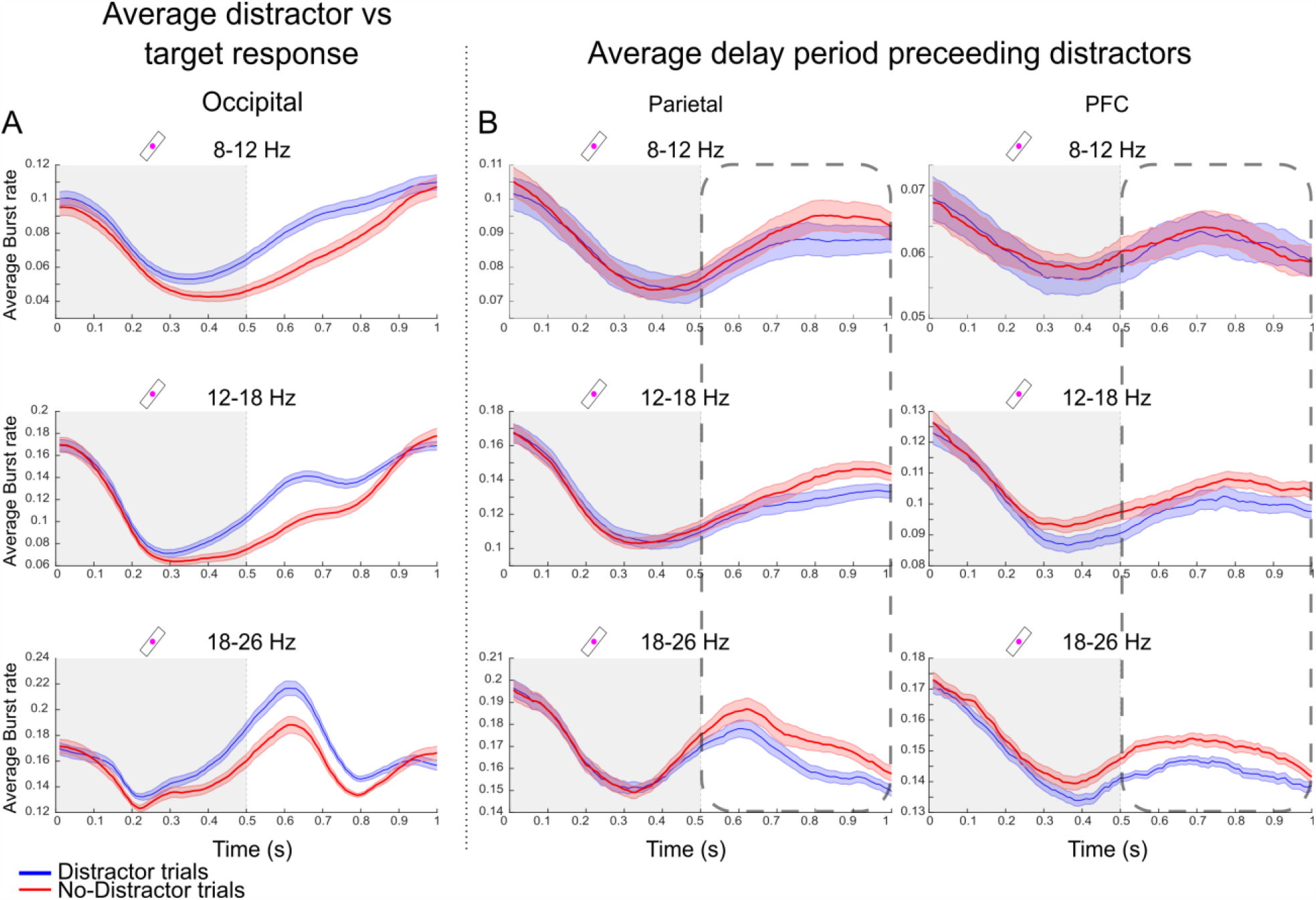
Bursting around distractor presentations. A) Average burst rates (as shown in Figure 5A) computed over a 1 second period from the onset of bars number 2 and 3 for each frequency band and the occipital area. The grey shaded area indicates the presentation period of bars 2 and 3. B) Average burst rates (as shown in Figure 5A) computed over a 1 second period from the onset of bars number 1 and 2 for each frequency band and the parietal and PFC areas. The dotted box represents the delay period preceding the presentation of distractors (blue) or targets (red) for bar 2 and 3 in the sequence.

Second, in contrast to the above, there was *elevated* bursting in No-Distractor trials compared with Distractors trials in the delay periods *preceding* the distractors. This was primarily seen in parietal and prefrontal sensors and mostly in the beta bands (Figure 6; similar qualitative differences in alpha were not significant, see Figure 5). Thus, in the period before target items were about to be presented, there was elevated bursting in prefrontal and parietal regions. This is again consistent with the suggested inhibitory role, with elevated beta bursting proactively freeing up resources in higher order regions for the upcoming item by down-regulating information already held in WM.

Third, the gradual decrease in beta bursting over the course of the sequence was only observed over parietal and prefrontal, not occipital sensors (Figure 5B). Prefrontal sensors overall had similar behaviour as the parietal, with a key difference that there was also a brief period of elevated bursting after the second distractor consistent with removal of that information.

Fourth, in the memory delay following the sequence there was an elevation of bursts in all three frequency bands and all sensors. This difference likely reflected a difference in WM load (2 vs 4 items), and disappeared after the retro-cue, which effectively equalized the memory load between the two conditions.

Finally, the gradual decrease of beta, but not alpha, bursting over the course of the task was region specific. This gradient was observed in parietal and prefrontal but not occipital sensors (Figure 5B).

### Beta and alpha bursts correlated with encoding on single trials

The burst analysis offered an opportunity to establish beta and alpha bursts as a single-trial correlates of sensory filtering. To investigate this, we analysed how the amplitude of phase-locked power in the tagged frequencies during stimulus presentations were modulated by the presence of bursts (burst-triggered phase-locked power). For this we used all bursts from occipital sensors, i.e. the sensors in which tagged frequencies were modulated by distractors, occurring during the time of stimulus processing (between 200 to 700 ms from stimulus onset, the period in which tagged frequencies were elevated). Since both burst rates and phaselocked power showed strong modulation over the course of a stimulus presentation, burst-triggered power was then compared to surrogate data to test if the presence of bursts modulated power at the tagged frequencies (Methods). The surrogate data was produced by shuffling trial indexes (for each sensor, subject and condition independently) for which the associated phase-locked power was taken from. Thus, importantly, the temporal structure was preserved in surrogate data. This revealed that bursts in all three frequency bands had a small but significant inhibitory effect on phase-locked power (Figure 7). Again, there were some differences between bands. For alpha bursts, power was reduced following onset of bursts. For beta, there was in addition significantly elevated power prior to the onset of bursts. We interpreted this as beta bursts being more likely to occur when tagged frequencies around stimulus presentations happened to be elevated in power, consistent with a role in active regulation of processing.

**Figure 7.**
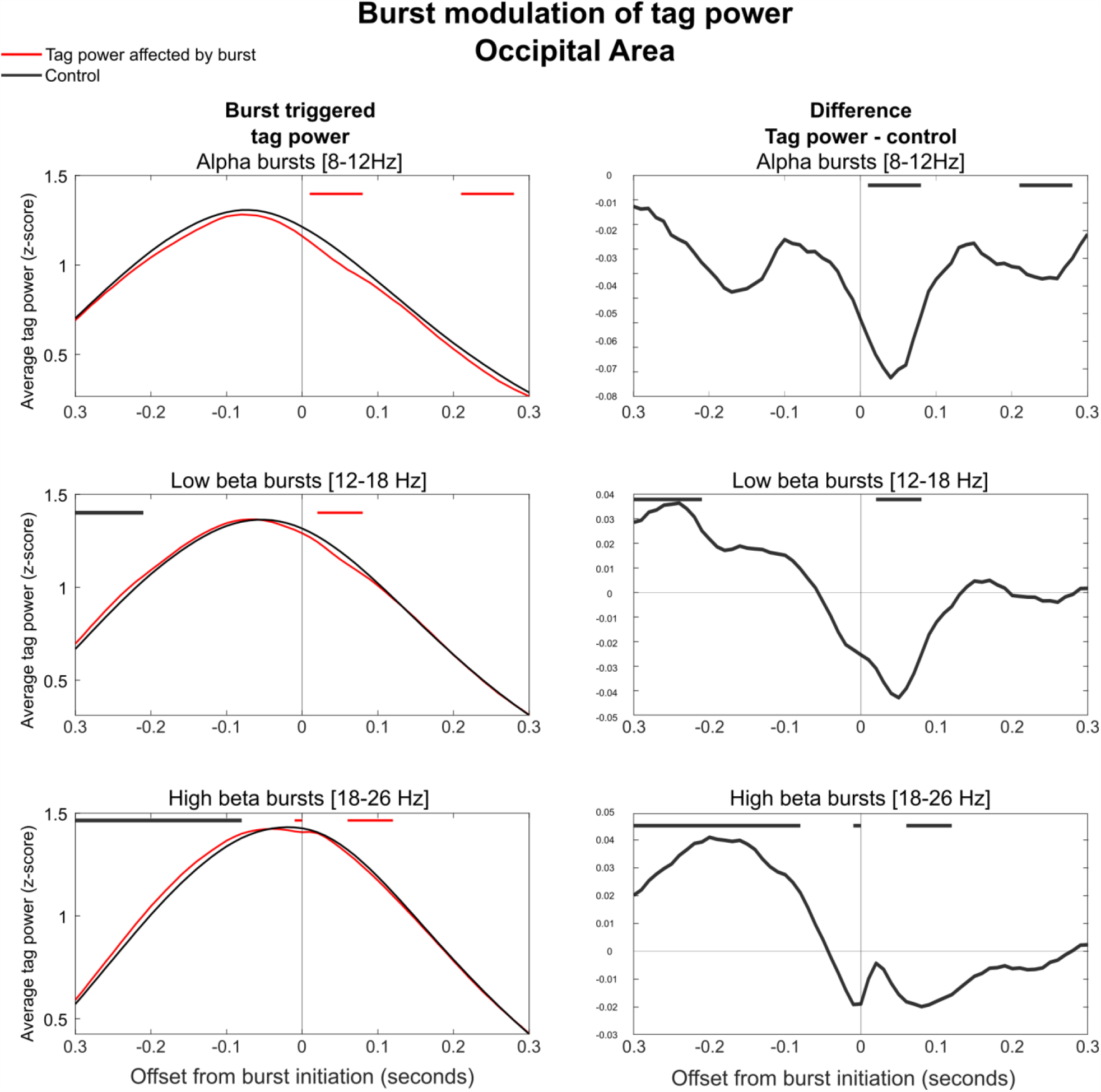
| Burst modulation of tag power for each frequency band. Left column shows the burst triggered average phase-locked power (z-score, see Methods) at the tagged frequency bands (red). Time 0 denotes the time of the detected burst onset. Displayed is also surrogate data, calculated by shuffling trial labels (within each condition and sensor independently) to estimate the power in the tagged frequencies drawn from the same distribution of times as the observed bursts (grey). Red bars denote periods in which power in the original data is lower than the surrogate data and grey bars indicate when power is higher in the original data than the surrogate data using two-sided permutation test at p < 0.001. The right column shows the difference between modulate tag power and surrogate data. The right column shows the difference between the actual data and the surrogate data.

## DISCUSSION

We set out to resolve the distinct neural correlates of WM control proposed by earlier work. Prior work has either suggested prefrontal beta bursts (non-human primates (Bastos et al., 2018; Lundqvist et al., 2016, 2018, 2023; Miller et al., 2018)) or occipital power fluctuations in alpha (humans (Ferrante et al., 2023; Gutteling et al., 2022; Popov et al., 2019; Riddle et al., 2020; Roux & Uhlhaas, 2014; Turner et al., 2023; Wolff et al., 2017; Zhigalov et al., 2019; Zhou et al., 2023)) as correlates of visual attention and WM control. We applied the single-trial burst analysis we developed for the non-human primate data (Lundqvist et al., 2016) on human data recorded with MEG. This demonstrated that WM-related beta bursts have a similar role in humans as in non-human primates. In addition, alpha bursting shared a similar behaviour to that of the beta bursts, but with some key differences. Significant differences between target and distractor processing were observed at different times and in different cortical regions for the two bands. Alpha bursts, primarily in occipital regions, suppressed distractor stimulus processing. Occipital beta bursts were instead elevated just as distractor stimuli were removed from the screen, in the transition from sensory processing to WM retention. In addition, beta bursts in parietal and prefrontal regions tracked sequence order and were elevated *before* relevant items. The latter pattern is consistent with inhibitory beta bursts in higher order cortex freeing up space in WM. Prior studies have indeed suggested that while alpha appears to be important in the filtering of irrelevant sensory information, it is not the full story and that several complementary mechanisms are at play (Jensen, 2023; Noonan et al., 2016).

Our existing non-human primate model suggests that the level of beta bursting reflects the level of inhibitory cognitive control (Lundqvist et al., 2016, 2018, 2023; Miller et al., 2018). This is based on simultaneous analysis of spiking and intracranial LFPs on the single trial level, where beta bursts suppress gamma bursts and spiking. It is also based on overall patterns of beta bursting during various cognitive operations, and how they differ on recording sites in which information is encoded into the patterns of spiking or not. Thus, in this model beta bursting is suppressed during encoding of information, and more so on recording sites where information is encoded, as well as during read-out of information. Beta bursting have intermediate levels when information is retained but not used, and then strongly elevated following each trial when information has to be cleared out, in particular on those sites information was encoded (Lundqvist et al., 2016, 2018; Miller et al., 2018).

Here we observed analogous beta and alpha burst rate patterns in occipital, parietal and prefrontal regions. To establish a single trial correlate of bursting using only non-invasive measures we used frequency tagging (Gutteling et al., 2022; Parkkonen et al., 2008; Zhigalov et al., 2019). It suggested indeed that both occipital beta and alpha bursting suppressed sensory processing on the single trial level. There was however an interesting difference between beta and alpha bursts in this regard. Processing tended to be elevated just prior to each beta burst, and not just suppressed during their presence. We interpret this as beta being up-regulated by a feedback mechanism when cortical levels of activity are too high. Alternatively, it was recently observed that beta bursts are preceded by a brief window of excitation before the longer lasting inhibition (Law et al., 2022). This could also explain the elevated levels of processing preceding beta bursts.

There were several other differences between alpha and beta bursting, and between the cortical regions. We argue these differences taken together suggest that alpha acts more as a sensory filter, while beta, particular in higher order areas, acts on information already within the cortex: Occipital alpha was suppressed more during the presentation of target items compared to distractors. The overall rates were not modulated by other task factors such as load or the passage of time within trials, however. Occipital beta bursts rates were also modulated by the presence of distractors in a similar way, but more focused around the time of the removal of the distractor from the screen (thus when it would have to be encoded into WM). Activity over parietal and prefrontal sensors had a quite different pattern with significantly elevated beta bursting *before* the presentations of target items (as opposed to upcoming distractors). We interpret this as down-regulation of existing WM-related activity, to reallocate resources for the upcoming target items in a proactive way. We draw these conclusions primarily based on timing of the alpha and beta bursting. Further experiments, where the timing of distractors and target items are varied, and in which distractors are not always predictable, may shed further light on this potential distinction between alpha and beta bursting.

In addition, beta (but not alpha) burst rates in parietal and prefrontal regions were strongly modulated by the passage of time within trials, with gradually lower burst rates as the trial advanced. Since this trend was equally strong in Distractor and No Distractor trials we find it likely to be tracking the passage of time and task structure, rather than being load dependent (which also changed with time but differently for the two types of trials). Thus beta bursting in higher order areas may be more directly linked to executive control functions, where keeping track of the various parts of a trial is essential. We have recently proposed that the spatio-temporal evolution of beta bursting patterns help implement cognitive operations by directing information flow to distinct patches of cortex during distinct parts of a task (Lundqvist et al., 2023). This shapes low-dimensional and task-related aspects of single neuronal spiking, that are needed to solve the cognitive task at hand (Badre et al., 2021; Xie et al., 2022).

The role of beta in WM executive function was also recently supported in clinical studies (Boo et al., 2023; Paulo et al., 2023). In patients with Parkinson’s disease insufficient cortico-striatal beta power suppression during encoding of information into WM was linked to diminished WM performance and correlated with symptom severity (Paulo et al., 2023). In obsessive compulsive disorder (OCD), the diminished prefrontal beta power rebound following trials was linked to the impairment in removal of information from WM (Boo et al., 2023).

Human studies have suggested that modulation of alpha power reflects filtering of sensory processing in attention and WM tasks (Sauseng et al. 2005; Popov et al. 2017; Zhou et al. 2023; Zhigalov et al. 2019; Roux and Uhlhaas 2014; Gutteling et al. 2022). Intracranial recordings demonstrate that alpha activity is finely tuned spatially, consistent with a role of selective suppression of unwanted information in a visual scene (Popov et al., 2019; Yuasa et al., 2023). Alpha power has also been reported to suppress distractors in different ways depending on the sensory modality in a WM task with visual and auditory inputs (Zhou et al., 2023). Thus we propose that whole-cortex burst patterns in alpha (primarily in sensory and parietal regions) and beta (primarily in parietal and prefrontal regions) dynamically evolve to orchestrate the flow of sensory information to be stored in or deleted from WM according to behavioural (task) demands. These cortical burst patterns are in turn likely coordinated by interactions with thalamus and basal ganglia (de Mooij−van Malsen et al., 2023; Jayachandran et al., 2023; Ketz et al., 2015; Law et al., 2022; Miller et al., 2018; Paulo et al., 2023; Sherman et al., 2016).

In sum, our results help unite the body of findings regarding the role of beta and alpha in attention and WM in different cortical areas, across human and non-human primates. They appear to serve distinct roles, that may be further teased apart by future experiments.

## METHODS

### Participants

We recruited 19 participants, 12 males and 7 females, aged 21-41 years (mean 27.1), with no known cognitive impairments and tested with Ishihara’s test for colour deficiency without remarks. The participants were primarily recruited students and received 500 SEK in compensation vouchers. One male did not perform the experiment due to metal interfering with the MEG scanner, and one male was excluded due to poor data quality resulting from excessive movement during the experiment resulting in a total of 17 participants. The Swedish Ethical Review Authority (Dnr 2021-00336) approved the study. All participants were thoroughly briefed and provided written consent to their participation.

### Experimental paradigm

The purpose of the task was to study burst dynamics across regions and frequencies during the presentation of stimuli and the effect of distractors. The task was a sequential working memory task with 4 foveally presented, randomly oriented bars (Figure 1). Before the onset of each sequence there was a pre-cue indicating if the current trial was a Distractor (50%) or No-Distractor trial (50%). Distractor and No-Distractor trials were randomly interleaved with the criterion that a maximum of 3 trials of a certain type could be presented in a row. On No-Distractor trials the participants were tasked with remembering the orientation of all four bars. On Distractor trials only the first and the fourth bars were to be remembered, and the second and third bars were distractors. The No-Distractor trials were indicated by a pre-cue with four parallel vertical solid bars, while the Distractor trials were indicated by the same bars except that the second and third bars were illustrated with dashed edges informing the participant that these were distractors (not to be remembered). Except for the precue, the two types of trials were visually identical, which allowed us to study the neural mechanisms of encoding, filtering and removing information by comparing a to-be memorized item with a distractor. Each trial commenced on a black background, with a white fixation dot during 500 ms, followed by a pre-cue shown for 500 ms. Next, four bars with random orientation, each covering a visual angle of 7.1º were shown sequentially for 500 ms, separated by a brief delay period of 500ms with a centrally placed white fixation dot. Each individual bar in the sequence was marked with a uniquely coloured fixation dot in the centre (the colour was randomly selected on each trial from red, green, blue and magenta). Following the presentation of the fourth bar, a white fixation dot was shown for 750 ms. The fixation thereafter changed colour (red, green, blue or magenta) acting as a retro-cue. The retro-cue identified the upcoming target for 750 ms after which the subject was presented with a randomly rotated probe. Using a 4-button control pad, the subjects had 5 seconds to rotate the probe to match the target. The subjects had to perform 400 trials in blocks of 40 with pauses in between. In each pause the subject was asked to decide when to continue, and in total the task took between 60-70 minutes for the subjects to complete.

### Frequency Tagging

Frequency tagging involves manipulating the spectral power. This is achieved by oscillatory modulation of the intensity of the stimuli, which entrains neural activity to the frequency of the modulation. This provides insights into when and where the sensory stimuli are processed in the brain (as measured by MEG). Frequency tagging of the stimuli has been applied in both auditory (Bharadwaj et al., 2014; Manting et al., 2023) and visual perception (Parkkonen et al., 2008; Zhigalov et al., 2019). We modulated the luminance of the cue and the four bars by a sine function phase-locked to the onset of the stimuli with tagging frequencies of either 31.1 or 37.1 Hz. We chose prime frequencies to avoid sub-harmonics, and the decimal was added from the result of a limited pilot study. Within each trial, the objects in the sequence were tagged at alternating frequencies (31.1 Hz and 37.1 Hz), with 50% of the trials tagging the first object with 31.1 Hz and the rest at 37.1 Hz. The stimuli with a frequency tag appeared as faint flickering and no visual difference between the two frequencies was seen or reported.

### Procedure, Materials and Data acquisition

Participants were sent information about the experiment in advance and upon arrival they were greeted and fully informed about the task. They signed a consent form, a MEG screening form and provided a suitable set of clothing to change into. All jewellery, hairpins and any other objects were removed at this stage. The participants were then prepared for the procedure by fitting electrodes and head digitalization. Next, they were led into a magnetically shielded room where the experiment was conducted. Before the MEG recordings were made, each subject was given time to practice the task for about 25-40 trials.

The MEG scanner was an Elekta Neuroma TRIUX 306-channel, located inside a 2-layer magnetically shielded room (www.natmeg.com). The procedure was presented on a FL35 LED DLP Projector from Projection Design running at 32 bit colour in 1920*1080 at 120 fps, and the experiment was performed using Presentation® software (Version 23.0, Neurobehavioural Systems, Inc., Berkeley, CA, www.neurobs.com). Eye movements and eye-blinks were recorded with an Eyelink 1000 binocular tracker from SR Research.

Data were recorded at the sampling rate of 1000 Hz for all 306 MEG channels, eye tracker channels, 16 event code channels and 2 channels dedicated to electrooculography and electrocardiography. Subjects submitted their responses using a 4-button inline pad from Current Design.

### Behavioural analysis

To analyse behaviour, we collected all responses and compared their angles to the target angles. The circular error, defined as the angle of the response subtracted from the angle of the target in a circular reference frame (from –90° to +90°), was calculated for all trials. We then took the absolute value of the error and applied a transformation, to assure that the assumptions of the general linear model were met. First all zero errors were set to 0.5 degrees. After that, we applied a Box-Cox transformation with parameter L=0.101. The parameter L was estimated to meet as close as possible the normality assumptions.

Posterior inspection of the residuals showed a good fit with the assumptions of normality (Q-Q plots) and homogeneity of the variances.

Mixed effects models where fitted in R (R Core Team (2023), n.d.). The following packages were used: lme4 (Bates et al., 2015), lmerTest (Kuznetsova et al., 2017), Mass (Venables & Ripley, 2002), performance (Lü-decke et al., 2021).

### Data preprocessing and frequency domain analysis

The recorded data files were run through Maxfilter software by Elekta Neuromag, with temporal signal space separation. Further analysis was done using Fieldtrip. Data files were segmented into trials starting one second before the trial start and one second after the submission of the response. Trials were demeaned, line noise removed and trials with jump artefacts, identified as a z-score greater than 80, were removed. This resulted in 15 trials removed on average per subject. Across the MEG channels 60 ICA components were calculated using FastICA and then used to automatically identify and remove EOG and muscle artefacts using procedures provided by Fieldtrip. ECG artefacts were identified and removed using a semi-automatic procedure as described by Fieldtrip. The data was down sampled to 250 Hz.

Time frequency calculations were done for single trials in the range 5-45 Hz using adaptive, multiplicative superlets (Moca et al., 2021) with order ranging from 1 to 10, and a base wavelet length of three cycles. Only data from planar gradiometers were used. For analysis where a baseline was applied, a trial-by-trial z-score baseline was calculated based on the 250 ms epoch before the precue onset. Total power was calculated from the pre-processed activity data, and Induced power was calculated by removing the ERF from the activity data and applying the same method. Phase-locked power was calculated by subtracting Induced power from Total power (Cohen, 2014).

### Burst extraction

To identify bursts of high power, we first specified three frequency bands of interest: 8-12 Hz (alpha), 12-18 Hz (low beta), or 18-26 Hz (high beta). The cut-off at 26hz was chosen to avoid spectral leakage from the tagging frequency at 31.1hz. Within each band and for each sensor we calculated the temporal profile of the induced power using adaptive superlets. Bursts were defined as intervals within each trial where instantaneous power was above 1.5 standard deviations above the mean (running average of the last 10 full trials), and with the duration of at least two cycles of the average frequency of the band (Lundqvist et al., 2016). For each subject, the burst events were averaged across the stimuli into a burst rate per subject, from which a grand average burst rate (per frequency band and cortical region) was calculated. Individual burst times were also kept for further single trial analysis.

### Burst triggered frequency tagged power

Burst events for each frequency band, subject, channel and trial were extracted for a window of 200-700ms after the onset of a frequency tagged stimuli. Each event was associated with its descriptive information (subject, channel, trial type), and the spectral power within the tagged frequency (narrow band of 4 Hz) was extracted from phase-locked power, during a window of 30ms before and after the onset of the burst event. The burst triggered frequency tagged phase-locked power was calculated by averaging events within subject, and then across subjects.

### Statistical information

Permutation tests (Maris & Oostenveld, 2007) were performed for the burst activity across subjects.

To relate bursts rates between the tasks (Figure 5 and 6), modulation indices were calculated and the following steps were done

Trial types were shuffled within each channel and subject in order to preserve the structure of the data.

A grand average was calculated across subjects and channels, and compared to modulation indices of the actual data averaged across subjects.

The points above were repeated 1000 times, and 99.9% percentile limits were identified.

The test for burst frequency tagged power required a different approach (Figure 7). A matrix containing all burst events with their tagging power and descriptive information was constructed, and trial numbers were shuffled 1000 times within their descriptive groups. Averages were calculated and compared to actual data in order to calculate p-values, and tests were performed at the 99.9% percentile.

## ACKNOWLEDGEMENT

This work was funded by ERC starting grant 949131, and Swedish Research Council (Vetenskapsrådet) grants 2018-04197, 2022-02328.

## References

Badre, D., Bhandari, A., Keglovits, H., & Kikumoto, A. (2021). The dimensionality of neural representations for control. Current Opinion in Behavioral Sciences, 38, 20–28. 10.1016/j.cobeha.2020.07.002

Bastos, A. M., Loonis, R., Kornblith, S., Lundqvist, M., & Miller, E. K. (2018). Laminar recordings in frontal cortex suggest distinct layers for maintenance and control of working memory. Proceedings of the National Academy of Sciences of the United States of America, 115(5), 1117–1122. 10.1073/pnas.1710323115

Bates, D., Mächler, M., Bolker, B., & Walker, S. (2015). Fitting linear mixed-effects models using lme4. Journal of Statistical Software, 67(1), 1–48. 10.18637/jss.v067.i01

Bharadwaj, H. M., Lee, A. K. C., & Shinn-Cunningham, B. G. (2014). Measuring auditory selective attention using frequency tagging. Frontiers in Integrative Neuroscience, 8, 6. 10.3389/fnint.2014.00006

Boo, Y. J., Kim, D.-W., Park, J. Y., Kim, B. S., Chang, J. W., Kang, J. I., & Kim, S. J. (2023). Altered prefrontal beta oscillatory activity during removal of information from working memory in obsessive-compulsive disorder. BMC Psychiatry, 23(1), 645. 10.1186/s12888-023-05149-1

Chatham, C. H., & Badre, D. (2015). Multiple gates on working memory. Current Opinion in Behavioral Sciences, 1, 23–31. 10.1016/j.cobeha.2014.08.001

Cohen, M. X. (2014). Analyzing neural time series data: theory and practice. The MIT Press. 10.7551/mitpress/9609.001.0001

D’Esposito, M., & Postle, B. R. (2015). The cognitive neuroscience of working memory. Annual Review of Psychology, 66, 115–142. 10.1146/annurev-psych-010814-015031

de Mooij-van Malsen, J. G., Röhrdanz, N., Buschhoff, A.-S., Schiffelholz, T., Sigurdsson, T., & Wulff, P. (2023). Task-specific oscillatory synchronization of prefrontal cortex, nucleus reuniens, and hippocampus during working memory. IScience, 26(9), 107532. 10.1016/j.isci.2023.107532

Ferrante, O., Zhigalov, A., Hickey, C., & Jensen, O. (2023). Statistical learning of distractor suppression downregulates prestimulus neural excitability in early visual cortex. The Journal of Neuroscience, 43(12), 2190–2198. 10.1523/JNEUROSCI.1703-22.2022

Goldman-Rakic, P. S. (1995). Cellular basis of working memory. Neuron, 14(3), 477–485. 10.1016/0896-6273(95)90304-6

Gutteling, T. P., Sillekens, L., Lavie, N., & Jensen, O. (2022). Alpha oscillations reflect suppression of distractors with increased perceptual load. Progress in Neurobiology, 214, 102285. 10.1016/j.pneurobio.2022.102285

Haegens, S., Luther, L., & Jensen, O. (2012). Somatosensory anticipatory alpha activity increases to suppress distracting input. Journal of Cognitive Neuroscience, 24(3), 677–685. 10.1162/jocn_a_00164

Hipp, J. F., Hawellek, D. J., Corbetta, M., Siegel, M., & Engel, A. K. (2012). Large-scale cortical correlation structure of spontaneous oscillatory activity. Nature Neuroscience, 15(6), 884–890. 10.1038/nn.3101

Jayachandran, M., Viena, T. D., Garcia, A., Veliz, A. V., Leyva, S., Roldan, V., Vertes, R. P., & Allen, T. A. (2023). Nucleus reuniens transiently synchronizes memory networks at beta frequencies. Nature Communications, 14(1), 4326. 10.1038/s41467-023-40044-z

Jensen, O. (2023). Gating by alpha band inhibition revised: a case for a secondary control mechanism. PsyArXiv.

Ketz, N. A., Jensen, O., & O’Reilly, R. C. (2015). Thalamic pathways underlying prefrontal cortex-medial temporal lobe oscillatory interactions. Trends in Neurosciences, 38(1), 3–12. 10.1016/j.tins.2014.09.007

Kuznetsova, A., Brockhoff, P. B., & Christensen, R. H. B. (2017). lmertest package: tests in linear mixed effects models. Journal of Statistical Software, 82(13), 1–26. 10.18637/jss.v082.i13

Law, R. G., Pugliese, S., Shin, H., Sliva, D. D., Lee, S., Neymotin, S., Moore, C., & Jones, S. R. (2022). Thalamocortical Mechanisms Regulating the Relationship between Transient Beta Events and Human Tactile Perception. Cerebral Cortex, 32(4), 668–688. 10.1093/cercor/bhab221

Liesefeld, A. M., Liesefeld, H. R., & Zimmer, H. D. (2014). Intercommunication between pre-frontal and posterior brain regions for protecting visual working memory from distractor interference. Psychological Science, 25(2), 325–333. 10.1177/0956797613501170

Lüdecke, D., Ben-Shachar, M., Patil, I., Waggoner, P., & Makowski, D. (2021). performance: An R Package for Assessment, Comparison and Testing of Statistical Models. The Journal of Open Source Software, 6(60), 3139. 10.21105/joss.03139

Lundqvist, M., Bastos, A. M., & Miller, E. K. (2020). Preservation and Changes in Oscillatory Dynamics across the Cortical Hierarchy. Journal of Cognitive Neuroscience, 32(10), 2024–2035. 10.1162/jocn_a_01600

Lundqvist, M., Brincat, S. L., Rose, J., Warden, M. R., Buschman, T. J., Miller, E. K., & Herman, P. (2023). Working memory control dynamics follow principles of spatial computing. Nature Communications, 14(1), 1429. 10.1038/s41467-023-36555-4

Lundqvist, M., Herman, P., Warden, M. R., Brincat, S. L., & Miller, E. K. (2018). Gamma and beta bursts during working memory readout suggest roles in its volitional control. Nature Communications, 9(1), 394. 10.1038/s41467-017-02791-8

Lundqvist, M., Rose, J., Herman, P., Brincat, S. L., Buschman, T. J., & Miller, E. K. (2016). Gamma and beta bursts underlie working memory. Neuron, 90(1), 152–164. 10.1016/j.neuron.2016.02.028

Maris, E., & Oostenveld, R. (2007). Nonparametric statistical testing of EEG- and MEG-data. Journal of Neuroscience Methods, 164(1), 177–190. 10.1016/j.jneumeth.2007.03.024

Ma, W. J., Husain, M., & Bays, P. M. (2014). Changing concepts of working memory. Nature Neuroscience, 17(3), 347–356. 10.1038/nn.3655

Manting, C. L., Gulyas, B., Ullén, F., & Lundqvist, D. (2023). Steady-state responses to concurrent melodies: source distribution, top-down, and bottom-up attention. Cerebral Cortex, 33(6), 3053–3066. 10.1093/cercor/bhac260

Miller, E. K., Lundqvist, M., & Bastos, A. M. (2018). Working Memory 2.0. Neuron, 100(2), 463–475. 10.1016/j.neuron.2018.09.023

Moca, V. V., Bârzan, H., Nagy-Dăbâcan, A., & Mureşan, R. C. (2021). Time-frequency super-resolution with superlets. Nature Communications, 12(1), 337. 10.1038/s41467-020-20539-9

Noonan, M. P., Adamian, N., Pike, A., Printzlau, F., Crittenden, B. M., & Stokes, M. G. (2016). Distinct mechanisms for distractor suppression and target facilitation. The Journal of Neuroscience, 36(6), 1797–1807. 10.1523/JNEUROSCI.2133-15.2016

Parkkonen, L., Andersson, J., Hämäläinen, M., & Hari, R. (2008). Early visual brain areas reflect the percept of an ambiguous scene. Proceedings of the National Academy of Sciences of the United States of America, 105(51), 20500–20504. 10.1073/pnas.0810966105

Paulo, D. L., Qian, H., Subramanian, D., Johnson, G. W., Zhao, Z., Hett, K., Kang, H., Chris Kao, C., Roy, N., Summers, J. E., Claassen, D. O., Dhima, K., & Bick, S. K. (2023). Corticostriatal beta oscillation changes associated with cognitive function in Parkinson’s disease. Brain: A Journal of Neurology, 146(9), 3662–3675. 10.1093/brain/awad206

Popov, T., Gips, B., Kastner, S., & Jensen, O. (2019). Spatial specificity of alpha oscillations in the human visual system. Human Brain Mapping, 40(15), 4432–4440. 10.1002/hbm.24712

Popov, T., Kastner, S., & Jensen, O. (2017). FEF-Controlled Alpha Delay Activity Precedes Stimulus-Induced Gamma-Band Activity in Visual Cortex. The Journal of Neuroscience, 37(15), 4117–4127. 10.1523/JNEUROSCI.3015-16.2017

Riddle, J., Scimeca, J. M., Cellier, D., Dhanani, S., & D’Esposito, M. (2020). Causal evidence for a role of theta and alpha oscillations in the control of working memory. Current Biology, 30(9), 1748-1754.e4. 10.1016/j.cub.2020.02.065

Rosanova, M., Sarasso, S., Massimini, M., & Casarotto, S. (2020). Cortical Excitability, Plasticity and Oscillations in Major Psychiatric Disorders: A Neuronavigated TMS-EEG Based Approach. In B. Dell’Osso & G. Di Lorenzo (Eds.), Non invasive brain stimulation in psychiatry and clinical neurosciences (pp. 209–222). Springer International Publishing. 10.1007/978-3-030-43356-7_15

Roux, F., & Uhlhaas, P. J. (2014). Working memory and neural oscillations: α-γ versus θ-γ codes for distinct WM information? Trends in Cognitive Sciences, 18(1), 16–25. 10.1016/j.tics.2013.10.010

R Core Team (2023). (n.d.). R: A language and environment for statistical computing. R Foundation for Statistical Computing, Vienna, Austri. Retrieved November 5, 2023, from https://www.R-project.org

Sauseng, P., Klimesch, W., Stadler, W., Schabus, M., Doppelmayr, M., Hanslmayr, S., Gruber, W. R., & Birbaumer, N. (2005). A shift of visual spatial attention is selectively associated with human EEG alpha activity. The European Journal of Neuroscience, 22(11), 2917–2926. 10.1111/j.1460-9568.2005.04482.x

Sherman, M. A., Lee, S., Law, R., Haegens, S., Thorn, C. A., Hämäläinen, M. S., Moore, C. I., & Jones, S. R. (2016). Neural mechanisms of transient neocortical beta rhythms: Converging evidence from humans, computational modeling, monkeys, and mice. Proceedings of the National Academy of Sciences of the United States of America, 113(33), E4885–94. 10.1073/pnas.1604135113

Turner, W., Blom, T., & Hogendoorn, H. (2023). Visual Information Is Predictively Encoded in Occipital Alpha/Low-Beta Oscillations. The Journal of Neuroscience, 43(30), 5537–5545. 10.1523/JNEUROSCI.0135-23.2023

Venables, W. N., & Ripley, B. D. (2002). Modern Applied Statistics with S (4th ed.). Springer New York. 10.1007/978-0-387-21706-2

Vogel, E. K., McCollough, A. W., & Machizawa, M. G. (2005). Neural measures reveal individual differences in controlling access to working memory. Nature, 438(7067), 500–503. 10.1038/nature04171

Wolff, M. J., Jochim, J., Akyürek, E. G., & Stokes, M. G. (2017). Dynamic hidden states underlying working-memory-guided behavior. Nature Neuroscience, 20(6), 864–871. 10.1038/nn.4546

Xie, Y., Hu, P., Li, J., Chen, J., Song, W., Wang, X.-J., Yang, T., Dehaene, S., Tang, S., Min, B., & Wang, L. (2022). Geometry of sequence working memory in macaque prefrontal cortex. Science, 375(6581), 632–639. 10.1126/science.abm0204

Yuasa, K., Groen, I. I. A., Piantoni, G., Montenegro, S., Flinker, A., Devore, S., Devinsky, O., Doyle, W., Dugan, P., Friedman, D., Ramsey, N., Petridou, N., & Winawer, J. (2023). Precise spatial tuning of visually driven alpha oscillations in human visuaul cortex. BioRxiv. 10.1101/2023.02.11.528137

Zhigalov, A., Herring, J. D., Herpers, J., Bergmann, T. O., & Jensen, O. (2019). Probing cortical excitability using rapid frequency tagging. Neuroimage, 195, 59–66. 10.1016/j.neuroimage.2019.03.056

Zhou, Y. J., Ramchandran, A., & Haegens, S. (2023). Alpha oscillations protect working memory against distracters in a modality-specific way. Neuroimage, 278, 120290. 10.1016/j.neuroimage.2023.120290

